# A reassessment of positive growth effects of expressed random sequence clones in *E. coli*

**DOI:** 10.64898/2026.04.08.717174

**Authors:** Sven Künzel, Carla Borish, Cornelia Burghardt, Corinna Heidinger, Diethard Tautz

## Abstract

*De novo* gene emergence from non-coding sequences is increasingly recognized as an important evolutionary mechanism, yet the functional potential of random sequences remains debated. Previous experiments suggested that expression of random sequence clones in *Escherichia coli* can enhance growth of the cells bearing them, i.e. they provide a fitness advantage. However, these findings have been questioned, regarding potential confounding effects of the clone mixtures and a possibly negatively acting peptide expressed from the cloning vector. Here we performed controlled competitive growth assays using a defined subset of 64 random sequence clones representing a spectrum of fitness effects. Experiments across multiple conditions, including two different growth cycle durations, induction states, and replicate sets, showed high technical reproducibility and consistent clone-specific growth trajectories for the majority of the clones, but for some also influences of genomic background and experimental conditions. While vector-derived constructs that inhibit the vector-coded peptide expression showed the same fitness improvements relative to the parental vector that were previously shown, several random sequence clones exhibited higher positive selection coefficients under conditions of exponential growth. These effects persisted even when negative clones were excluded, indicating that they are not driven by competition dynamics with negative clones. Our results demonstrate that positive growth effects of random sequence clones cannot be explained by clone mixture and vector artifacts alone. Instead, a subset of random sequences confers genuine fitness advantages comparable to beneficial mutations observed in experimental evolution studies. These findings provide strong experimental support for the capacity of random sequences to generate adaptive functions and underscore their role in *de novo* gene evolution.

**Significance statement:** This study provides robust experimental evidence that a subset of random DNA sequences can confer genuine fitness advantages in Escherichia coli, independent of previously proposed artifacts such as vector effects or clone competition. Based on controlled competitive assays across multiple conditions, the results show that these adaptive effects are reproducible and comparable to beneficial mutations observed in experimental evolution. These findings strengthen the case that random sequences can serve as a meaningful source of functional innovation, supporting their role in de novo gene evolution.

## Introduction

Comparative genomic analyses have shown that genes can be created *de novo* from previously non-coding sequences in the genome (reviewed in (Tautz and Domazet-Loso 2011; Andersson, et al. 2015; Van Oss and Carvunis 2019; Weisman 2022; Zhao, et al. 2024; Casola, et al. 2025; Xia, et al. 2025; Bornberg-Bauer and Eicholt 2026)). While this mechanism has long been thought to be rare (Tautz 2014), there are now many well-studied cases from all domains of life that suggest that the *de novo* emergence of genes from previously non-coding sequences may be a rather frequent process. To better understand how easy it is to generate genetic functions out of random sequences, there have been multiple experiments based on the expression of more or less randomly synthesized DNA sequences that were tested for function in different experimental regimes. One set of experiments aimed to rescue genetic defects (e.g. (Digianantonio and Hecht 2016; Babina, et al. 2023; Frumkin and Laub 2023), to convey resistance to chemical treatments or antibiotics (e.g. (Hoegler and Hecht 2016; Knopp, et al. 2019; Knopp, et al. 2021)) or resistance to virus infection (Frumkin, et al. 2025) by random sequence constructs. Another set of experiments has focused on tracing competitive growth differences in pools of clones (Bao, et al. 2017; Neme, et al. 2017; Castro and Tautz 2021; Bhave and Tautz 2022; Aldrovandi, et al. 2024) triggered by the expression of random sequences.

While all of these experiments have demonstrated the potential of generating genetic function out of random sequences, there remains an uncertainty of how important this mechanism is for evolutionary processes. At least for the experiments that aimed for genetic rescue through random sequences, it was shown that the expression of the random sequence in a wildtype background impedes cell growth, or is at least neutral, i.e. a fixation of such sequences would be expected to be rare under natural conditions when the selection trigger is not present. On the other hand, our first experiments to measure effects of random sequences on growth of wildtype cells under competitive conditions showed that up to 25% of the analyzed clones could enhance the growth of cells (Neme, et al. 2017). This result seemed to be rather surprising and it was therefore met with skepticism (Weisman and Eddy 2017). There were two main arguments against this experimental approach. First, the enrichment of clones in cycles of competitive growth of a whole library does not per se imply a positive growth effect, but could be a passive byproduct of interaction with other cells, or a general composition shift due to declining numbers of clones with strong negative effects. Second, if the vector used to express the random sequences would itself have a negative effect on the growth of the cells, this could mean that the cloned sequences in this vector simply relieve this negative effect, i.e. a true growth advantage is not proven. This latter point was experimentally addressed by a small set of experiments by (Knopp and Andersson 2018) from which the authors concluded that there is no evidence for a beneficial fitness effect of random sequences. While their experiments had indeed suggested a vector effect, the magnitude of this effect was small and would not have been enough to explain the strong effects that we found for a subset of the clones (Tautz and Neme 2018). Still, the issue was not fully resolved, which caused subsequent authors to dismiss our initial results as having an experimental design problem (Knopp, et al. 2021; Frumkin and Laub 2023).

However, the experiments have less a design problem, than a genetic verification problem. Rescuing a known phenotype allows a straight forward genetic verification, but it is much more difficult to trace the genetic basis for a general growth advantage, since any pathway could be affected. One approach is to do an RNASeq analysis with the clones that show a growth advantage, but this did not lead to a clear pathway identification either (Bhave and Tautz 2022). Hence, competitive growth experiments remain the best possible design for the general question of the relative frequency of beneficial effects from random sequences when one can control for a possible vector effect. We have therefore used here the same general design, but specifically tuned towards determining experimental variability parameters and using the vector and derived vector constructs (Knopp and Andersson 2018) as reference points in a controlled mix of clones. We find that the technical repeatability of the experimental design is very good, while the measured growth trajectories for a subset of random sequence clones can be influenced by the experimental parameters and that genomic background mutations in the host cells can also play a role. Still, under conditions of exponential growth, we can reproduce the differential vector effects shown by (Knopp and Andersson 2018) but show at the same time that these cannot explain the positive effects on growth that we a see for a subset of the random sequence clones. Further we show that this pattern is consistent when one removes all negative clones from the experiment, i.e. it is not due to a competition effect between clones. We consider the results from these experiments, in combination with previous re-analyses (Castro and Tautz 2021; Bhave and Tautz 2022) as evidence that the dismissal of our original results was not justified.

## Results

In a new set of clone competition experiments, we have retrieved a subset of 64 random sequence clones (suppl Table S1) from the original library representing a range from clones found to be positive, neutral or negative in previous analyses (Neme, et al. 2017; Castro and Tautz 2021).

Overnight cultures from these clones were mixed in approximately equal amounts for generating a sub-library for the new competitive growth cycle experiments. Using a defined subset of clones allowed to follow each individual clone in its growth trajectories across the cycles, rather than having them on a background of many low frequency clones in the original library.

We used two different cycle times, 24h-cycles that result in growth saturation for each cycle and 3h-cycles that keep the growth within the exponential phase. Each experiment was run in five technical replicates, without and with IPTG induction each (minus versus plus in the notations). We obtained samples from each cycle to determine the frequency changes of the clones by short read sequencing of the inserts. The numbers of reads for each clone were counted to serve as measure for their frequency in the whole population of clones.

We did a total of five separate experiments with the full clone set (expA to expE), two with five replicates each and three with two sets of five replicates in parallel (designated with 1 and 2 respectively), amounting to a total of eight datasets for minus and plus IPTG condition each (Table 1). In an additional experiment we tested a subset of positive clones only in two sets of five replicates each (expF). Raw counts as derived from the fastq files are provided in suppl. File S2, normalized counts in suppl. File S3.

**Table 1:** Growth cycle experiments performed.

| experiment | cycle time | number of cycles | clone status |
| --- | --- | --- | --- |
| expA | 24 hours | 4 | clones directly from library |
| expB | 24 hours | 4 | retransformed clones + vector constructs |
| expC1 | 24 hours | 4 | retransformed clones + vector constructs |
| expC2 | 24 hours | 4 | retransformed clones + vector constructs |
| expD1 | 3 hours | 4 | retransformed clones + vector constructs |
| expD2 | 3 hours | 4 | retransformed clones + vector constructs |
| expE1 | 3 hours | 3 | retransformed clones + vector constructs |
| expE2 | 3 hours | 3 | retransformed clones + vector constructs |
| expF1 | 3 hours | 3 | subset of retransformed clones + vector constructs |
| expF2 | 3 hours | 3 | subset of retransformed clones + vector constructs |

### Variances in replicates

In a first step, we determined coefficient of variances (CV) of normalized counts between the five technical replicates of each experiment. These variances are relevant for assessing the replicability with the given experimental setup. Figure 1 shows the average CV values across all eight datasets. They are generally below 0.1 for the minus IPTG replicates (average 0.07) and below 0.2 for the plus IPTG replicates (average 0.11). Interestingly, they rise with cycle number, especially in the plus IPTG replicates. This is expected, since the replicates constitute separate growth trajectories that should diverge over time, especially when growth affecting clones are expressed. Hence, we can conclude from this analysis that the experiments are reliable, with the level of technical and/or biological noise being mostly much below 25%.

**Figure 1:**
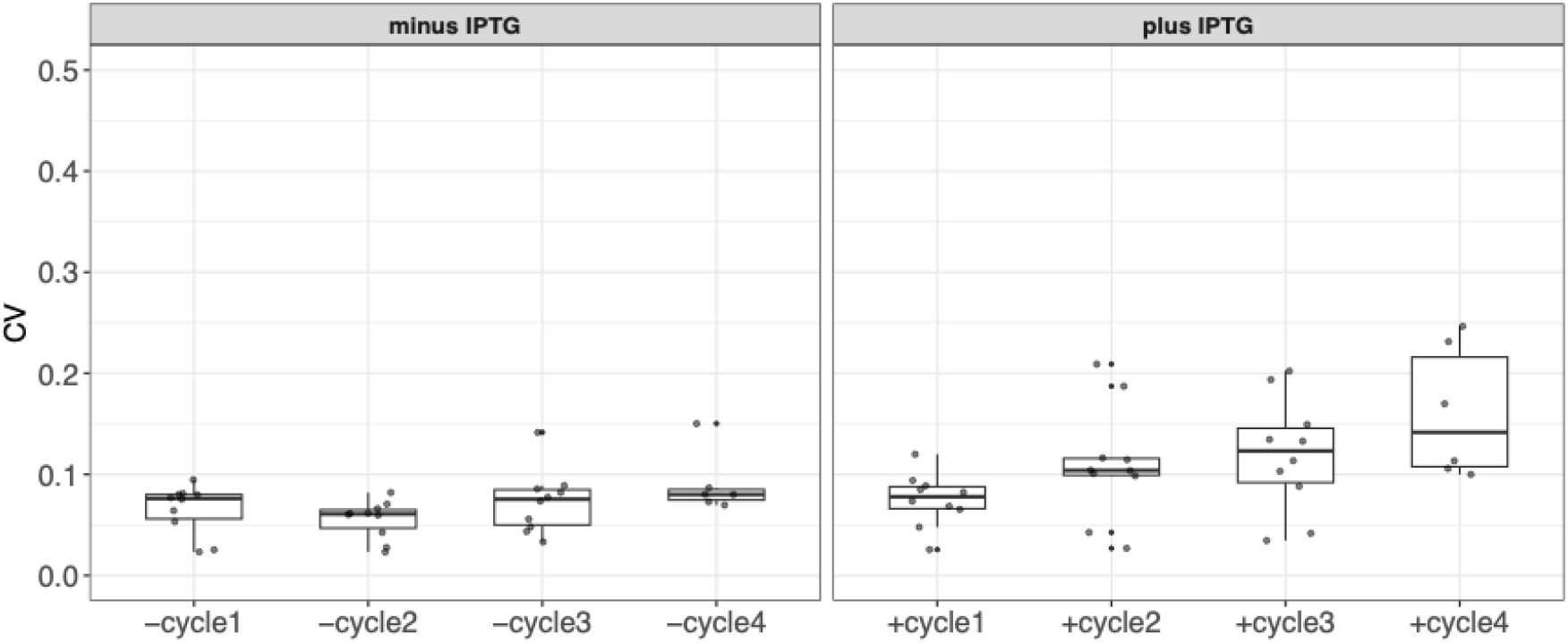
Average coefficients of variation (CV) for the five technical replicates across all experiments.

### Growth change effects

To quantify the growth changes induced by the random sequence clones, we used the linear trend function in DESeq2 to calculate selection coefficients per experimental cycle. We converted the DESeq2 linear slope to natural log, which makes it more directly comparable to the calculation of selection coefficients during exponential growth of bacterial cultures (DYKHUIZEN 1990), as it was also used in (Knopp and Andersson 2018). The results if the DESeq2 analysis for all experiments are listed in suppl. File S4, the distribution of the selection coefficient values across all experiments is summarized in Figure 2. While it is evident that the addition of IPTG has a profound effect on the growth trajectories of the clones, it is important to point out that some clones show also significant growth changes when no IPTG is added. This could be due to some leakage from the promotor as this effect is specifically evident for clones that show a strong negative effect under IPTG induction. But we note that the vector constructs show also a positive growth effect under minus IPTG conditions in the 3h cycles. It is unclear why this happens, but this indicates that the selection coefficients measured under the minus IPTG condition need to be subtracted from those of the plus IPTG condition to evaluate the specific effect of the random sequence induction.

**Figure 2:**
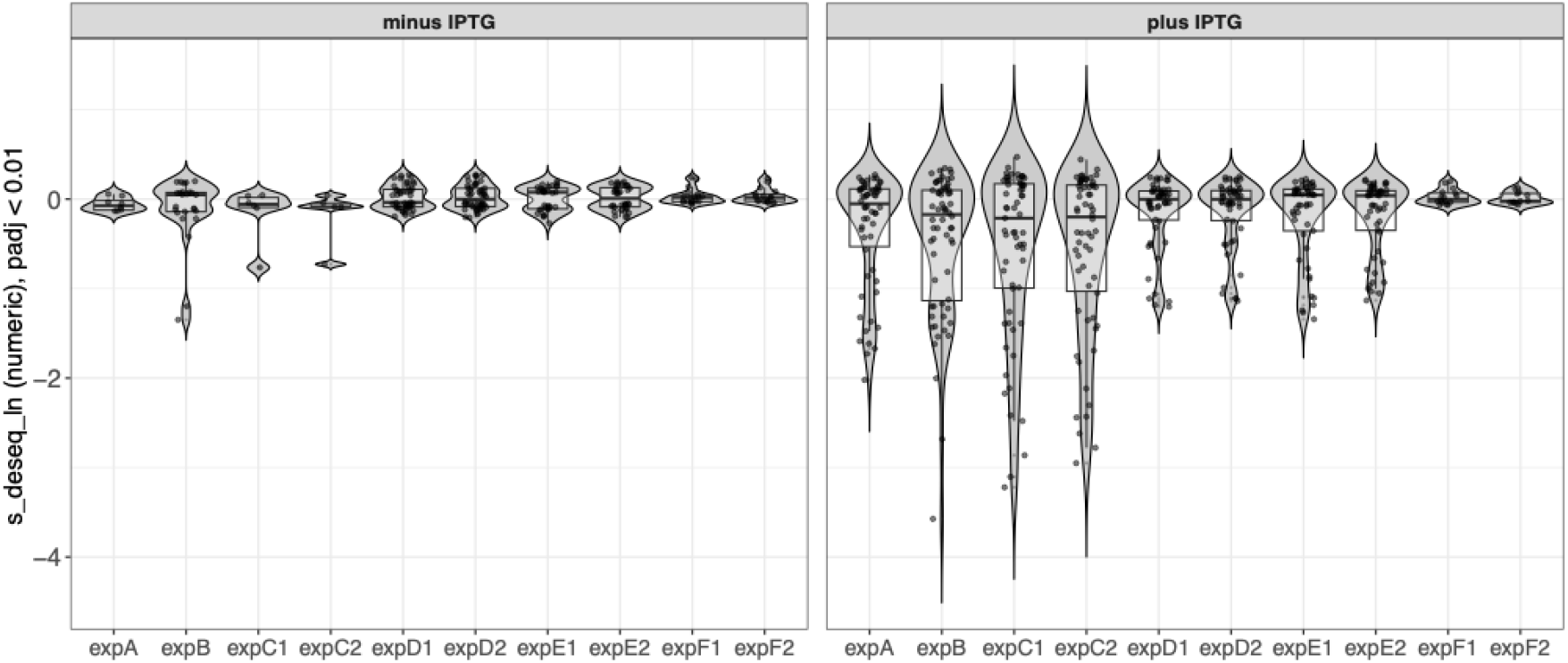
Violin plots of selection coefficients across all clones for all experiments. Note that only values that showed a significant change (Wald test, p < 0.01, with multiple test correction) are included in these plots. The full data are provided in suppl. File S4.

### Assessment of the influence of the genomic background

In a first experimental comparison, we asked whether the effects for each clone are indeed due to the plasmid or could be modified by background mutations in the bacterial host genome. For this purpose, we used the data from two experiments with four 24h cycles of growth, whereby the first included a starting pool of clones that were directly derived from the original random sequence library (expA), while the second was composed of clones that were re-derived through transforming the extracted plasmids from the original clones into new competent cells of the same bacterial host strain (expB). Comparison of the net selection coefficients for all clones shows a high correlation of 0.78 (Pearson correlation, p<<0.001) but also two outlier clones (Figure 3).

**Figure 3:**
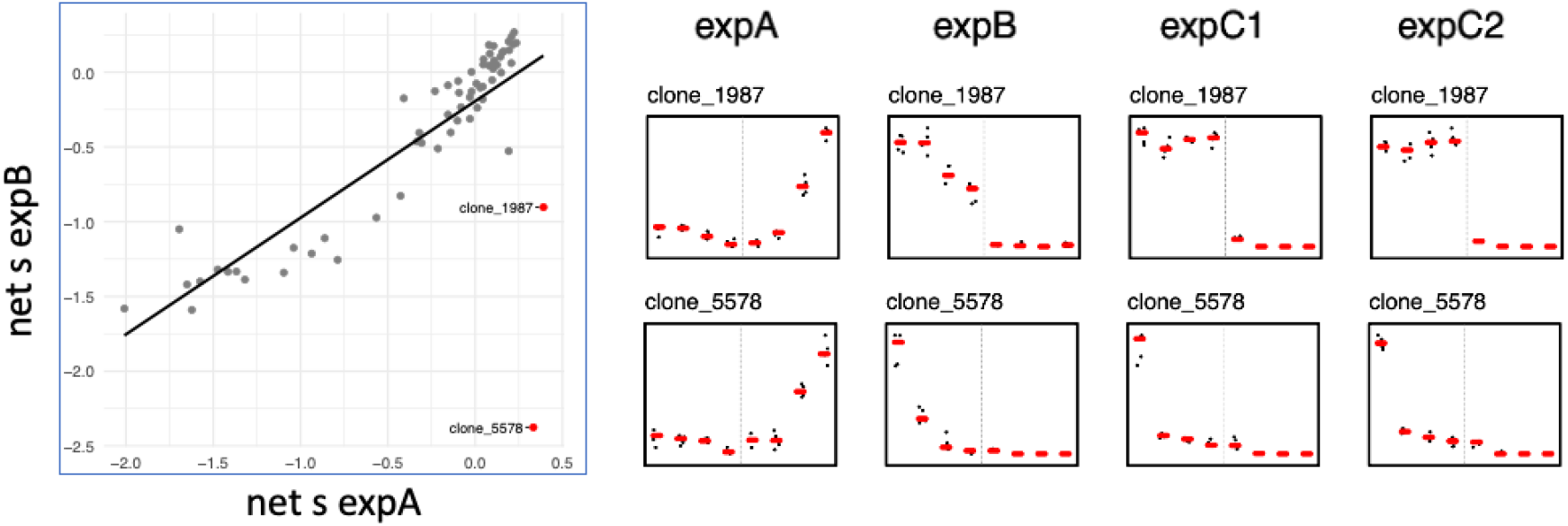
Comparison of clone selection coefficients (left panel) and growth trajectories of outlier clones (right panels) before and after transformation into new cells (expA versus expB). Outliers are defined by a Bland-Altman test with MAD threshold 4. The right panels show normalized counts for each of the four growth cycles as dots and their averages as red bars. The left halves of the plots represent the cycles without IPTG induction, the right halves the cycles plus IPTG induction.

For the two outlier clones (clone_1987 and clone_5578), we find a complete reversal from a positive growth trajectory towards a negative one. Given that the negative status is confirmed for these clones in expC (Figure 3, right), we conclude that there has been indeed a genome modification in the original clones that allowed them to be positive instead of negative. Interestingly, for clone_5578, the growth trajectory is negative even under the minus IPTG condition, implying a leakiness of the promotor in combination with a very strong negative effect.

From these results we conclude that genomic background of the cells can indeed influence the effects of the clones in the respective cells. Accordingly, for the further comparisons, we do not include the data from expA.

### Repeatability of overall trajectories

The experiments expC, expD and expE were conducted with two sets of five replicates each, run in parallel and analyzed independently. This tests for the repeatability of the calculation of the selection coefficients of the clones within a given experiment. Experiments expD and expE were done under short cycle conditions (i.e. 3h cycles) to compare repeatability with the long cycles (i.e. 24h) of expB and expC. Finally, in expD versus expE we compared results between 3 cycles and 4 cycles. Figure 4 shows the overall correlation plots for all of these experiments.

**Figure 4:**
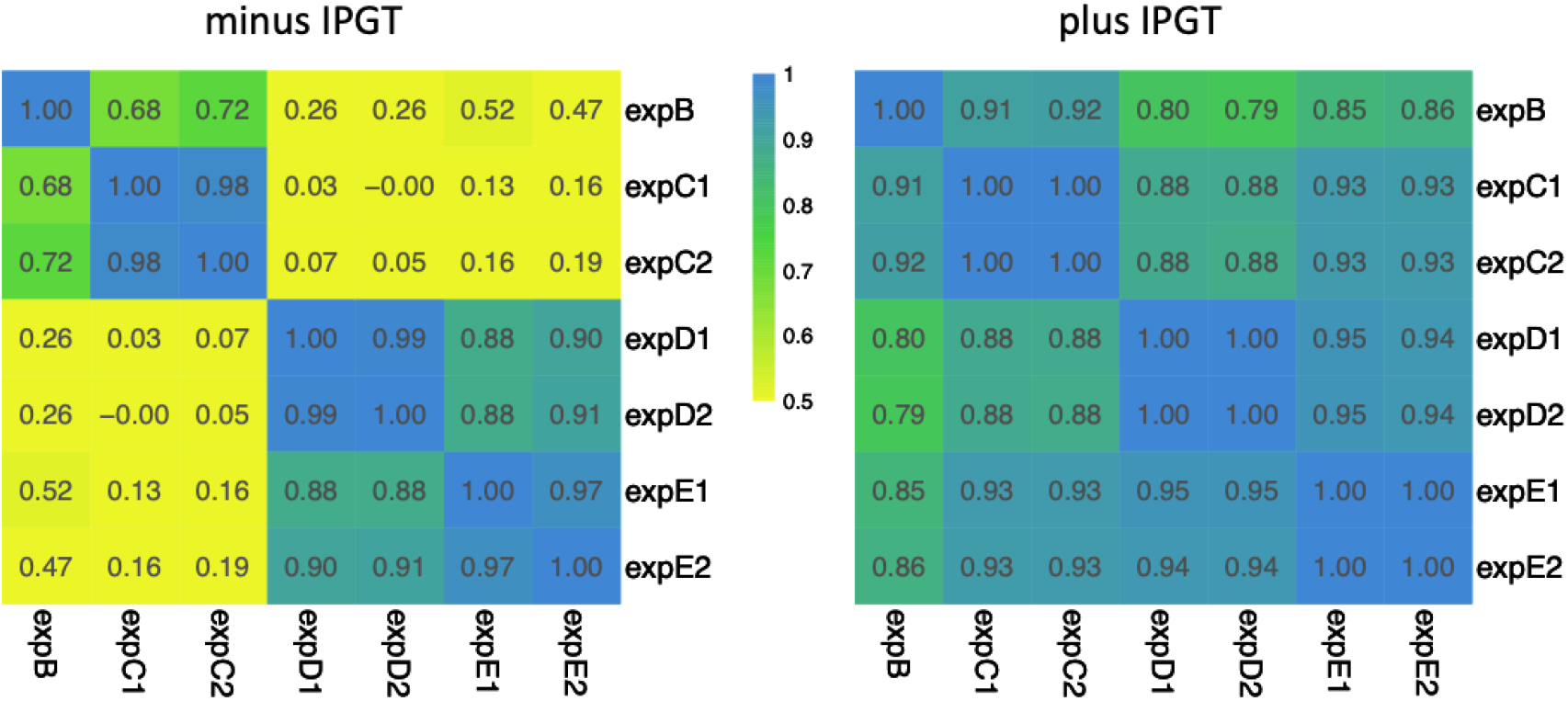
Correlation matrices for selection coefficients per cycle for the different experiments that include all clones, without (left) and with (right) IPTG induction. The inset numbers represent the correlation values. Numbers >0.4 are all significant (p < 0.01, Pearson correlation). The full data for the selection coefficients in all experiments are included in suppl. File S4.

For all three experiments conducted with 2×5 replicates, we find a very high correlation between the two sets of replicates, both for the minus IPTG and plus IPTG conditions, supporting the notion that the experimental procedure is robust, when the same conditions are applied. For the minus IPTG condition, the replicability between experiments was lower, more so for the 24h cycles (expB and expC), but still significant. Little correlation was found for the comparison between 24h cycles and 3h cycles in the minus IPTG condition.

In the plus IPTG condition, we found generally high correlations across all experiments, the highest for the 2×5 replicates, followed by the cycle times (24h - expB and expC, versus 3h - expD and expE). This suggests that there are some differences in clone performance that depend on the cycle times, although the overall replicability is high.

### Distribution of selection coefficients

As discussed in the introduction, the fundamental question to be addressed with these experiments is whether the positive clone effects could be simply ascribed to a vector effect. For this interpretation it is assumed that the RNA or the short peptide that is expressed from the vector wthout clone insert would excert a negative effect on cell growth. If this is replaced with an insert that is less negative, it could give the impression that it has a positive effect on cell growth.

To compare the overall vector and vector construct selection effects to all tested clones, we calculated the net selection coefficents, based on the 3h cycles, and ranked them in descending order (Figure 5). These cycles showed the highest replicability (see above) and are closest to the conditions where growth is measured during the exponential phase of cells, i.e. the conditions used in (Knopp and Andersson 2018). The plot shows that the vector constructs have higher net selection coefficients than the vector, confirming the general findings from (Knopp and Andersson 2018) on the relative performance of these clones. However, several of the positive clones have even higher selection coefficients, including two previously individually analysed clones from (Neme, et al. 2017) (clone 4 and clone 32). The third clone that was found positive in (Neme, et al. 2017), clone 600, was found neutral in the current experiments (Table 2), but this confirms also the test of this clone in (Knopp and Andersson 2018). We conclude therefore that our experiments are qualitatively comparable to the ones in (Knopp and Andersson 2018).

**Table 2.**
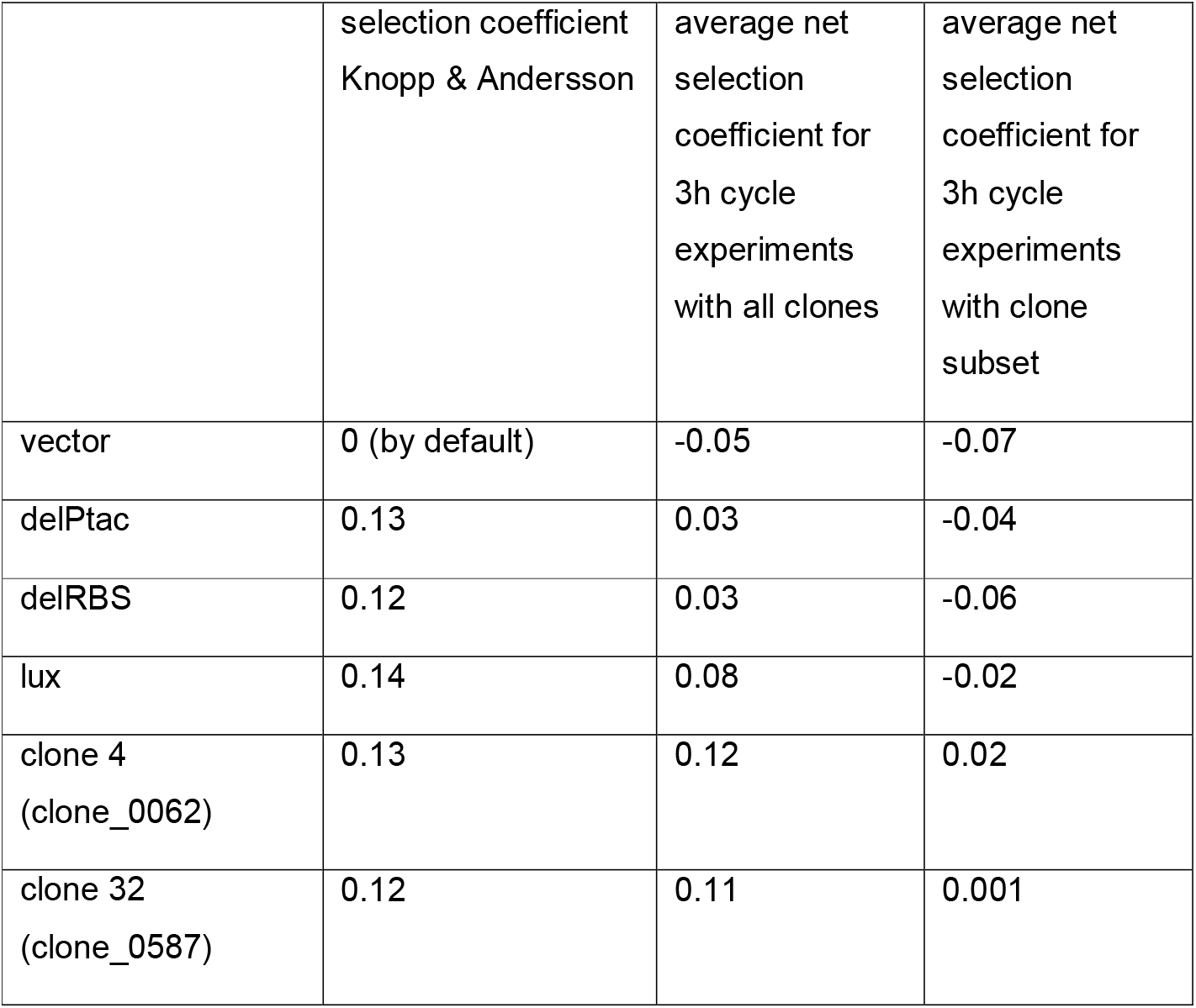
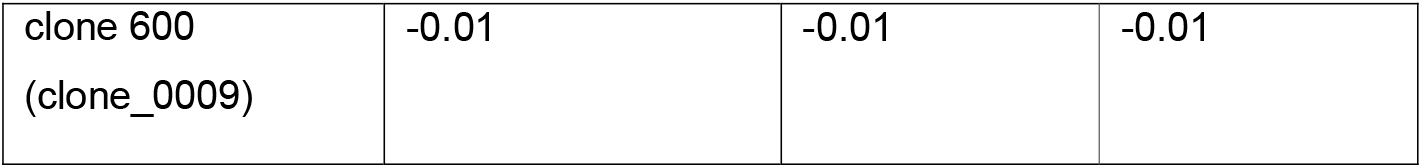
Comparison of selection coefficent measures for several clones in (Knopp and Andersson 2018) and in the present study.

**Figure 5:**
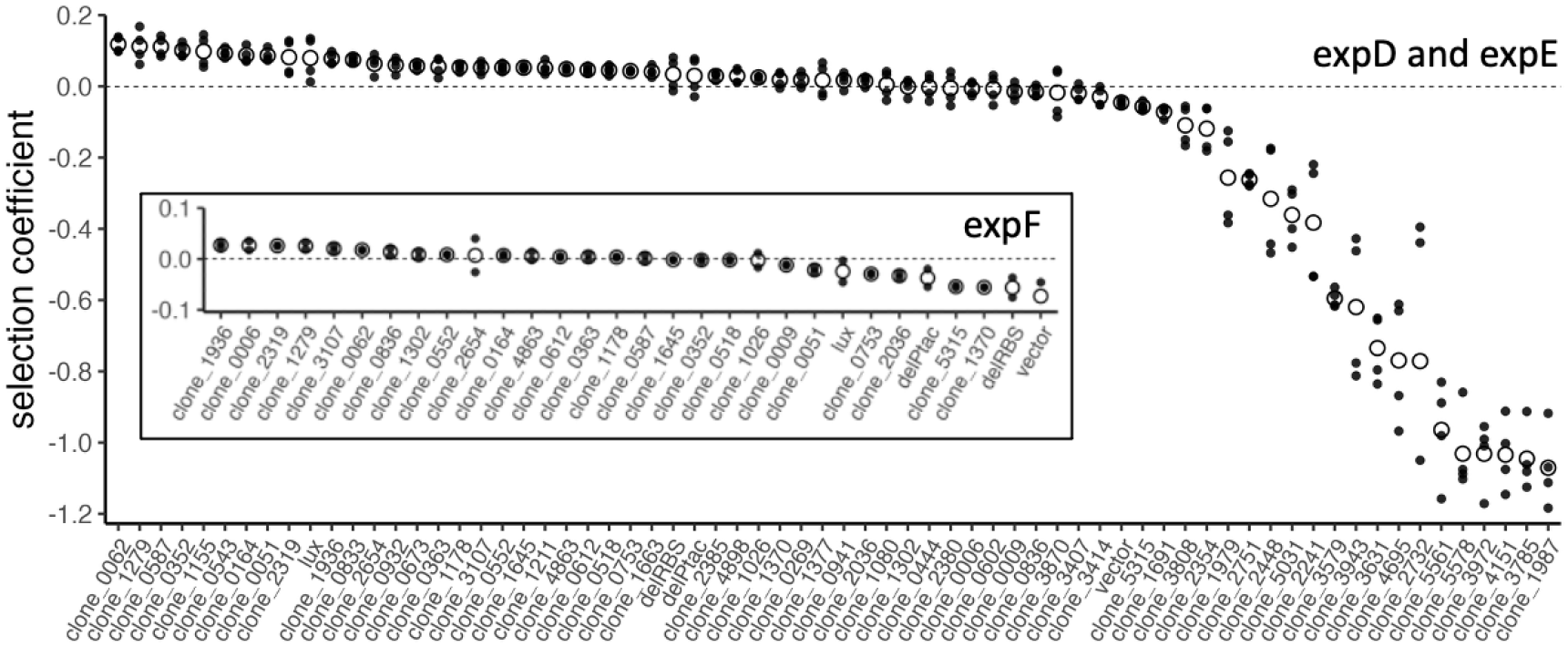
Net selection coefficients for all clones in the 3h cycles. Dots represent the actual values from the different experiments, circles the averages. Clone sorting is according to average values. The larger plot includes all clones from expD and expE, the inset the reduced set of clones from expF

### New clone mixture

Given that the experiment with all clones includes also many with a strong negative effect, we conducted an additional 3h cycle experiment with a subset of 26 clones that had shown higher selection coefficients than the vector, plus the four vector clones (expF1 and expF2). The results are shown in the inset of Figure 5. Again we see the vector having the lowest selection coefficient and the three vector constructs having higher ones (Table 2). However, many of the random sequence clones have still higher selection coefficients. Note that the overall selection coefficients are smaller in this experiment, presumably because the competetion between the clones is higher than in the experiment with the full set of clones.

## Discussion

The present study was specifically designed to assess whether the positive growth effects that are triggered by the expression of random sequences in individual clones could be due to an experimental design problem. For the study we used clones from previously published studies and conducted dedicated experiments to test for repeatability, vector effects and the influence of negative clones in the mixture.

With respect to repeatabilty of the individual clone effects, we find high consistency for experimental replicates (Figure 1) and within a given experimental regime, i.e. 24h cycles or 3h cycles, but somewhat less between the different regimes (Figure 4). Still, 37 out of the 68 clones showed consistently either a positive (N=17) or negative (N=20) net selection coefficient across all 7 experiments covering both experimental conditions (suppl. Table 1).

There was also a consistent effect of expression induction through the addition of IPTG. The spread of selection coefficients for individual clones was substantially larger under induced conditions for all experiments that included the full set of clones (Figure 2). However, we note that significant growth differences were also observable for some clones without IPTG induction. This suggests that carrying different versions of the plasmids can already have an effect on growth, possibly through some leakage of the promotor or DNA structure effects that influence the replicability of the plasmid. Hence, to assess the true selection differential that is generated by the induction of the random sequence expression, we calculated the net selection coefficients by subtraction of the s values of the minus IPTG measurements from the s values of the plus IPTG measurements.

The net selection coefficients ranged from -3.2 to 0.47 across all experiments. The average selection coefficients for the subset of clones with fully consistent positive growth effects ranged from 0.05 to 0.21. However, we noted that the net selction coefficient spread is smaller when only the clones were competed that showed positive selection coefficient in the full mixture experiments (expF - Figure 5). The maximum net selection coefficient was in this case 0.04. These values can be compared with the Lenski long-term evolution experiment (LTEE), where selection coefficients for beneficial mutations ranged between 0.006 to 0.059 (Imhof and Schlötterer 2001). Hence, we seem to capture a similar range of selection coefficients in our experiments.

Despite the high repeatability between experiments, we found also some clones that were more variable with respect to their growth dynamics between experiments. For two of the clones, we identified the genomic background as a source for causing this variability, since retransformation into a new background changed their growth trajectories from positive to negative (Figure 3). For the other clones with inconsistent trajectories, we assume that slight variations in experimental conditions, such as slight fluctuations in incubation conditions or growth medium composition might cause this. Such a condition-dependent effect on competitive growth was also found for the random sequence antitoxin RamF when it was expressed in cells without the toxin (Frumkin and Laub 2023).

### Vector and vector constructs

The relative growth effects of the vector and the vector constructs that were studied individually in (Knopp and Andersson 2018) were very well reproducible in the mixture experiments in the pool of clones. The vector expresses a peptide of 37aa and (Knopp and Andersson 2018) had suggested that this could be slightly deleterious. We find average net selection coefficients between -0.07 to 0.04 for the vector in our experiments (Table 2). The three vector constructs were designed to suppress the expression or translation of this peptide, with the expectation that the corresponding clones should perform better than the vector in the competitive growth condition. This is indeed the case throughout all of our experiments, whereby the removal of the promotor (delPtac) and the removal of the ribosome binding site (delRBS) have similar effects on growth, while the insertion of the premature termination signal (lux) has an even higher growth advantage (Table 2), both in the (Knopp and Andersson 2018) experiments, as well as in our experiments. These results imply that it does not make a major qualitative difference of whether the clones are tested in a pool or in isolation, but experimental conditions can influence the strength of the effects. This is particularly clear for expF which excluded the negative clones, such that the remaining positive clones would compete against each other. In this case we find on average lower selection coefficients for each clone, and even negative ones for the vector constructs, but still a similar relative order for their effects (Figure 5).

The assumption that the positive clones merely relief the negative effects of the vector peptide would imply that the maximum selection coefficient of the clones should not exceed the selection coefficient measured for the lux vector construct, since this introduces a transcription terminator that should abolish any expression. However, at least for the 3h experiments this is not the case. Here we find that multiple clones exceed the s measured for lux (Figure 5).

### Conclusions

Our experiments show that competition experiments in clone mixtures are a faithful method to measure relative fitness for clones expressing random sequences after IPTG induction. However, we found that it is necessary to correct the selection coefficients with those measured in a parallel setup without IPTG induction, since clones can show variable growth trajectories even without overexpressing the cloned sequence. At least for the condition under exponential growth of the clone mixtures, we find clones that have higher positive selection coefficients than a reference vector clone that does express a peptide from the respective promotor. Hence, we conclude that random sequence clones can indeed convey a true fitness benefit to their host cells. A recent analysis of predicted microproteins in the range of sizes as our random sequence peptides in bacteria has shown that they are abundant and can have fitness benefits for their host cells. Comparative analysis has shown that many of them have emerged *de novo* out of intergenic, i.e. presumably random, sequences (Fesenko, et al. 2025). Hence, our findings with synthesized random sequences bear also on the actual evolutionary processes that contribute to *de novo* gene formation.

## Methods

### Clones

All clones were derived from a library of random sequences cloned into the pFLAG-CT expression vector (Sigma-Aldrich, catalogue no. E8408) according to the procedure described in (Neme, et al. 2017). The random sequence length was 150bp, corresponding to 50 amino acids when fully translated, but usually shorter due to premature stop codons (the actual length distributions of peptides in the library are described in (Castro and Tautz 2021)). The vector includes the strong Ptac promoter, regulated by the presence of the lacO sequences and inclusion of the lac repressor gene (lacI) on the plasmid. It drives a transcript that includes a ribosome binding site and a start codon, as well as a C-terminal FLAG sequence with a stop codon. The oligonucleotides including the randomized sequences was cloned between the HindIII and SalI sites of the multiple cloning site.

The library was plated in a dilution to randomly select individual clones. The cloned parts from these clones were amplified were amplified by PCR and their inserts were sequenced by Sanger sequencing. For each clone we recovered the information on its growth dynamics from the previously conducted bulk experiments (Castro and Tautz 2021). Based on these, we generated a mixture of 64 clones that were expected to have on average positive, negative or neutral effects on the growth of the cells (listed in suppl. Table S1). Note that the growth effects were not necessarily consistent between different previous experiments, hence the classification of clones for including them into the present experiment was considered to be preliminary. Still, we tried to cover the complete range from positive to negative. Note that we retained the original numeric nomenclature for the clones, but changed the prefixes from “BACT” to “clone”, e.g. “BACT000002380” is converted to “clone_2380” for easier readability.

To control for genomic background effects, we compared in a first experiment the clone performances of cells that came directly from the library, with performances of the same clones after plasmid extraction and retransformation. For the further experiments, we added three clones (delPtac, delRBS and lux) obtained from (Knopp and Andersson 2018) that were engineered to change regulatory components of the vector, as well as the vector itself. The delPtac vector construct was designed to test whether the assumed cost of the general expression from this promotor can be relieved by deleting the promotor. The delRBS removes the ribosome binding site and was constructed to test whether the RNA versus the translated peptide have a relevant activity on the growth of the cellls. The lux vector construct includes a premature transcription terminator sequence to abort the expression of either the RNA or protein from the vector. These clones were also newly re-transformed from plasmids and then added to the library stock. Information on these clones are added to the list provided in suppl. Table S1 and checked by sequencing the inserts. An alignment of the modified sequences for these vector constructs is provided in suppl. Figure S5.

### Clone competition experiments

All clones were grown individually in overnight cultures and were then mixed in equal proportions to generate the library for competition growth. The mixture was adjusted to 20% glycerol and frozen in aliquots at -70°. These stocks were used to initiate an experiment with growing them in an overnight culture in LB medium following the detailed descriptions in (Neme, et al. 2017; Castro and Tautz 2021). In short: 500 μL of the overnight culture were transferred into replicate 5 mL tubes containing 4.5 mL of LB medium with 100µg/ml ampicillin without or with 10^−3^ mol/L IPTG to induce expression of the random sequences. For each cycle, 500 μL of culture from each tube were used to seed a new replicate after over night of growth (37°C, 250 rpm). 3mL of each tube were used for plasmid extraction for sequencing.

We used two different experimental regimes, one with 4 cycles of 24h cultures, the other with 3 cycles of 3h each. Hence, the first regime assess effects at stationary growth pahse, the second in exponential growth phase. Each experiment included 5 replicates, some included two sets of five replicates run in exact parallels.

### Sequencing and read coverage extraction

Sequencing of the bulk plasmid preparations was done on a Illumina MiSeq sequencer, following the protocol provided in (Castro and Tautz 2021) using the MiSeq Reagent Kit v2 300-cycles. Single read with 251bp was sequenced. Read counts for each clone were extracted from the fastq_R1 reads using the function vcountPattern implemented in the Biostrings package in Bioconductor (Huber, et al. 2015; Pages, et al. 2025), based on matches with the 20nt search motif sequences for each clone listed in suppl. Table S1. We allowed up to one mismatch to account for sequencing errors. The search motifs are derived from the random sequence parts of the clones and they are therefore unique. For the three modified vector sequences we derived the search motifs from the regions where the sequences were manipulated (compare suppl. Figure S5). The search motif for the vector was derived from the part of the multiple cloning site that is removed in the random sequence clones.

However, since this part is not removed from two of the three manipulated vector clones, vector counts were corrected by subtracting the count values of delPtac and delRBS.

### Determination of growth trajectories and selection coefficients

To determine individual selection coefficients for each clone, we used a time-course differential expression analysis implemented in DESeq2 (Love, et al. 2014). We used the raw counts from each experiment and its replicates (suppl. File S2) as input and the different cycle samples as time points. For the plus IPTG analysis we included the raw counts from the first cycle of the minus IPTG replicates as 0 timepoint. The normalized read output from DESeq2 (suppl. File S3) was used to plot growth trajectories.

Selection coefficients were estimated using a generalized linear modeling framework implemented in the DESeq2 pipeline with time being treated as a continuous numeric variable.The negative binomial regression model estimates a log2 fold change per unit time, corresponding to the slope of abundance change across the time series. Model fitting was performed using the Wald test with local dispersion estimation. To improve effect size estimates, log2 fold changes were subsequently shrunk using the adaptive prior-based method implemented in the apeglm package. The resulting shrunken log2 fold change per unit time was converted to a natural log scale to obtain the selection coefficient. This value represents the instantaneous rate of change in log abundance per unit time and is reported in the output column s_seseq_ln in suppl. File S4. Thus, the selection coefficient reflects the exponential growth or decline rate of each clone, assuming approximately log-linear dynamics over time. Positive values indicate increasing abundance (positive selection), whereas negative values indicate decreasing abundance (negative selection).

## Data and script availability

The primary data (fastq files) and R-scripts used in this study are available at Edmond under the pre-view link: https://edmond.mpg.de/previewurl.xhtml?token=f1323ced-8a9c-45c6-a437-7b63c8ec9599. The processed data files are included in the suppl. files of the manuscript.

## Author contributions

DT and SK designed the experiments, SK, CB, CB and CH conducted the experiments, DT did the data analysis and wrote the manuscript.

## Acknowledgements

We thank Dan Andersson for providing the vector clone constructs. The work was supported by institutional funds of the Max-Planck Society to DT.

