## Supplementary figures and images for "A reassessment of positive growth effects of expressed random sequence clones in *E. coli*"

### supplemetal Figure S5

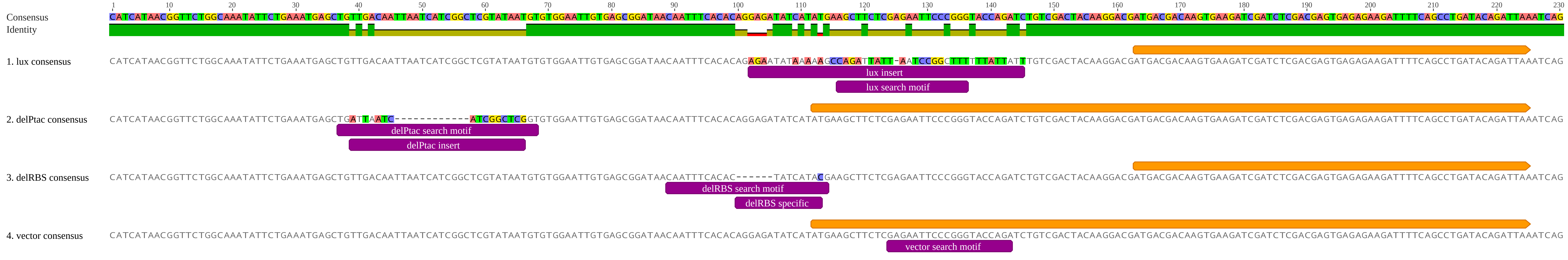
